# PCP and Septins govern the polarized organization of the actin cytoskeleton during convergent extension

**DOI:** 10.1101/2022.06.06.495000

**Authors:** Caitlin C Devitt, Shinuo Weng, Vidal D Bejar-Padilla, José Alvarado, John B Wallingford

## Abstract

Actin is generally required for cell movement but must be brought under the control of distinct developmental systems to drive specific morphogenetic processes, such as convergent extension (CE)^1^. This evolutionarily conserved collective cell movement elongates the body axis of all vertebrates and is governed by the Planar Cell Polarity (PCP) signaling system ^1,2^. Here, we sought to understand the interplay of PCP and actin during *Xenopus* CE. Our data provide new insights into the organization and emergence of polarity in this network and reveal new roles for the actin organizing Septins ^3^, thereby shedding light on the mechanisms by which developmental signaling systems exploit the ubiquitous machinery of cells to drive morphogenesis.

## Results and Discussion

Convergent extension requires the coordinated action of the PCP proteins ^1,2^ and core machinery of the actin cytoskeleton ^4–7^, but the relationships between these two elements remain incompletely understood. For example, the Septin proteins play key roles in actin assembly, organization, and dynamics ^3^ and are implicated in PCP and CE in frogs, mice, and fish ^6–10^. Septins appear to execute only a subset of PCP-dependent cell behaviors, as their loss recapitulates the severe tissue-level CE defects seen after core PCP disruption, yet leaves overt cell polarity intact ^6^. Moreover, several studies suggest a complex, reciprocal relationship between PCP and actomyosin during CE, with PCP signaling required to orient actomyosin contractions, and actomyosin also required for the polarized localization of PCP proteins ^11,12^.

Further obscuring our understanding of CE is the fact that cell movement generally requires the coordinated action of distinct but integrated populations of actin, such as the lamella and lamellipodia of migrating cells in culture ^13^ or the medial and junctional actin populations in cells engaged in apical constriction *in vivo* ^14,15^. In the context of *Xenopus* mesoderm CE, three such actin populations have been identified (Supp. Fig. 1), including a superficial, meshwork referred to as the “node-and-cable” system ^5,16–18^, a contractile network at deep cell-cell junctions ^7,19^, and mediolaterally oriented actin-rich protrusions which are present both superficially and deep ^5,19–21^. All of these actin systems are PCP dependent, but the precise relationships between PCP proteins, core actin regulators, and the polarization of actin networks remain poorly understood. Here, we exploited the amenability of the uniquely “two-dimensional” node and cable system to probe the relationship between PCP proteins, Septins, and the polarization of this actin network.

### PCP protein and Septin localization in the actin node and cable network during Xenopus convergent extension

We first examined localization of the core PCP proteins Vangl2 and Prickle1 in the node and cable system. During CE in zebrafish, both proteins were shown to localize to polarized foci, but the nature of these foci is unknown ^22–24^. We found that fluorescent fusions to either Vangl2 or Prickle1 in *Xenopus* were also present in foci, and moreover, that these foci were coincident with actin rich nodes (Fig. 1A, B). PCP foci (and thus actin-rich nodes, see Fig. 2, below) were not initially polarized, but became progressively biased to the anterior over time (Fig. 1D-E, H), suggesting that the PCP foci in *Xenopus* mesenchymal cells follow the pattern of progressive polarization observed for junction accumulation of PCP proteins in epithelial cells ^2,25^. The anterior bias of these foci was confirmed by immunostaining for endogenous Vangl2 (Supp. Fig. 2). By contrast, a fusion to the core PCP protein Dishevelled was localized to posteriorly biased foci that also developed progressively (Fig. 1C, G, F, H), similar to that reported in zebrafish ^22^, but these were *not* co-localized with actin-rich nodes.

**Figure 1:**
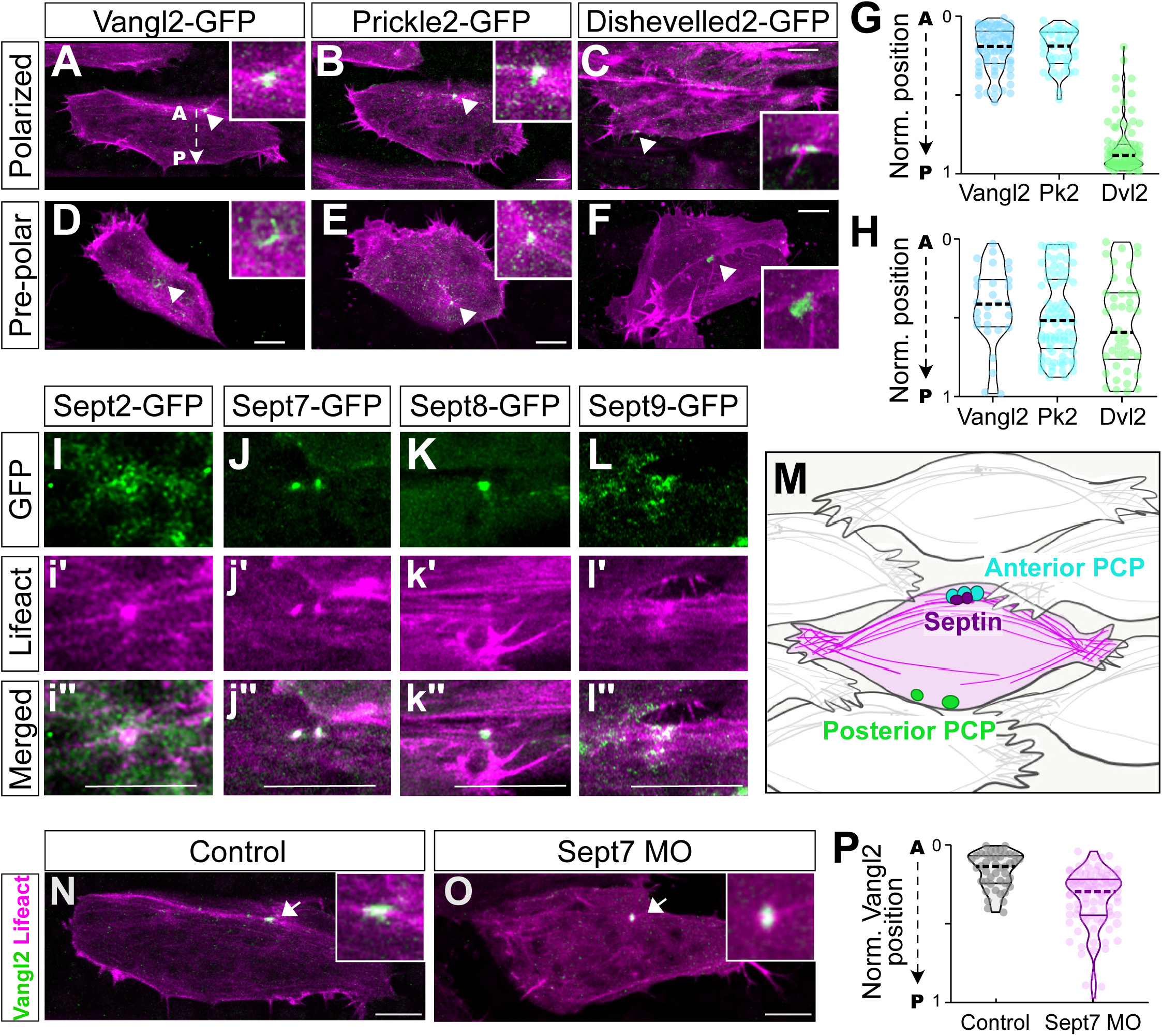
PCP protein and septin localization in actin rich nodes. **A-C**. Still images of indicated PCP proteins (green) and LifeAct (magenta) in polarized cells, insets show magnified view of nodes. **D-F.** Still images of indicated PCP proteins (green) and LifeAct (magenta) in pre-polarized cells, insets show magnified view of nodes. **G, H.** quantification polarization of PCP foci in polarized and pre-polarized cells. **I-L.** Images of indicated Septin subunit s(green) and LifeAct (Magenta) at nodes; Scalebar = 10 um. **M.** Schematic interpretation of data in A-L. **N, O.** Vangl2-GFP localization in control and Sept7MO cells. **P**. Quantification of AP positioning of Vangl2 puncta. N values and statistics are provided in Supplemental Information.

**Figure 2:**
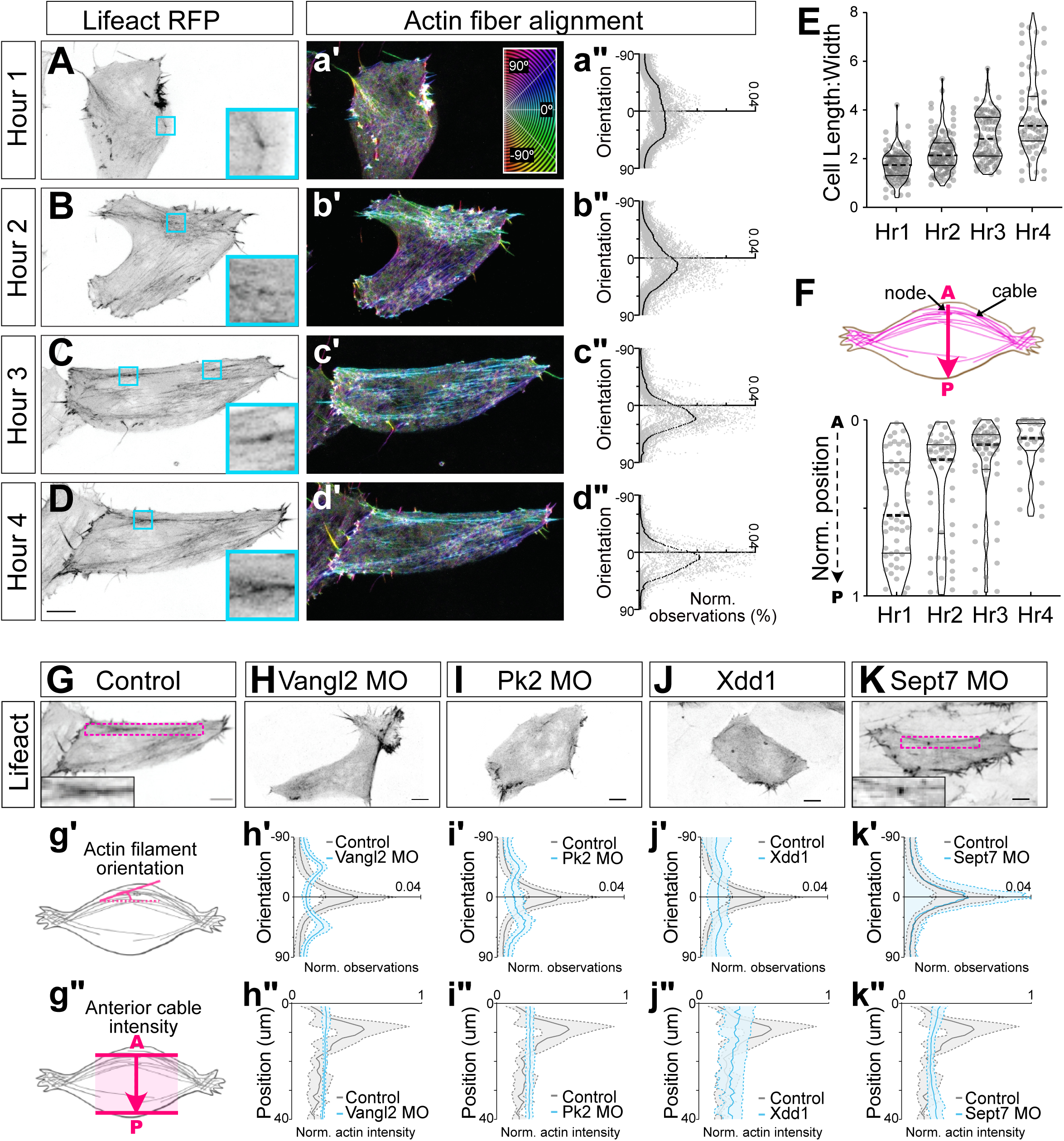
Progressive polarization of the actin node and cable system requires PCP and Septins. **A-D.** Individual cells mosaically labeled with LifeAct at times indicated at left (maximum intensity projections of cortical actin 10um into cell). **a’-d’.** Cyan box and inlay highlight individual nodes. Heatmap showing orientation of actin fibers quantified with OrientationJ (Legend in a’; cyan = mediolateral, red = anteroposterior). **a”-d”** Quantification of actin filament orientation. **E**. Quantification of cell length- to-width ratios over time. **F**. Upper panel, schematic of AP localization, arrow representing cellular anterior-posterior axis. Lower panel, quantification of node positioning over time. **G-K**. Representative images of cells with indicated manipulations. (Maximum intensity projection of cortical actin 10um into cell). Inlay shows actin node and cable, where present, indicated by magenta box). **g’, g”.** Schematics of quantification. **h’-k’:** Quantification of actin fiber alignment in cells. **h”-k”** Quantification of actin cable positioning. Scale= 10um. N values and statistics are provided in Supplemental Information.

We next assessed the localization of Septins, which are known to assemble into hetero-oligomeric complexes comprised of one each of four subgroups (see Supp. Fig. 3A, B)^26^. We first examined Sept2 and Sept7 subunits, which are essential for in CE ^6,7^. Both were enriched in anterior actin-rich nodes (Fig. 1I,J; Supp. Fig. 3D,E). To represent the two remaining Septin subgroups (Supp. Fig. 3A,B), we chose Sept8 and Sept9 based on their strong expression in the dorsal mesoderm during CE ^27^, and these, too, were strongly enriched at nodes (Fig. 1K, L; Supp. Fig. 3F,G). The localization of Septins reflected the diverse morphology of nodes, but as a population, their localization was highly consistent, as shown by overlapping peaks of Septin and actin intensity plots (Supp. Fig. 3d”-g”; Fig. 1M).

In light of their colocalization, we next probed the functional interrelationship between PCP proteins and Sept7. Interestingly, we found that actin rich, Vangl2+ nodes still form in the absence of Sept7, though they are mispositioned, and not strongly biased to anterior regions (Fig. 1N-P). Conversely, KD of Vangl2 elicited a robust and generalized loss of detectable of Sept7-GFP in cells (Supp. Fig. 4), which could reflect either disrupted localization/diffusion of Sept7 or a generalized loss of the protein. These interdependent phenotypes, with Vangl2 required for Sept7 localization and Sept7 also required for anterior positioning of Vangl2+ nodes, reflect the complex reciprocal relationship shown previously for PCP proteins and actomyosin during CE ^11,12^.

### A cryptic anteroposterior polarity in the node and cable system requires PCP proteins and Sept7

These data raise an interesting question, because while the orientation of actin fibers within the node and cable system and their myosin-driven contraction are polarized mediolaterally ^5,16–18^, no aspect of the system is yet known to be polarized in the A/P axis marked by PCP protein localization. These finding prompted us to further explore polarity within the node and cable system.

First, time-lapse analysis revealed that actin fibers were initially random, but became progressively aligned and oriented over time as cells progressively polarize and elongate ^20,28^(Fig. 2A-E). Interestingly, however, this progressive mediolateral polarization of fibers was coincident with the progressive polarization of actin-rich nodes to the anterior region of cells (Fig. 2F), reflecting the polarization of PCP proteins described above.

PCP disruption is known to completely abolish all aspects of planar polarity during CE, and accordingly, knockdown (KD) of Vangl2 or Prickle 2 (Pk2) or expression of dominant negative Disheveled (Xdd1), severely disrupted both mediolateral alignment of actin fibers (Fig. 2G, g’ – J, j’) and the anterior positioning of actin cables (Fig. 2g’’ – j’’). By contrast, Sept7 loss strongly disrupts CE at the tissue level but leaves overt cell polarity intact ^6^, and consistent with that effect, loss of Sept 7 *did not* affect the mediolaterally polarized alignment of individual actin filaments (Fig. 2K, k’) but *did* significantly disrupt the anterior enrichment of actin cables (Fig. 2k’’). This result was paradoxical. On the one hand, the lack of effect on actin fiber alignment was consistent with the normal overall cellular polarity in Sept7 KD cells ^6^. On the other, the effect on anterior actin cable positioning suggests the presence of a cryptic Sept7-dependent anteroposterior-oriented organization of the actin system in these cells.

This possibility prompted us to develop another metric of actin organization that was more granular than cable formation yet independent of actin fiber alignment. We therefore segmented LifeAct-GFP labeled actin fibers from our TIRF movies using SOAX software^51^ and used customized scripts to quantify their persistence length (L_p_)(Fig. 3A). This metric reflects the bending stiffness of polymers in isolation, but in the complex environment of the cell, we reasoned that it may serves instead as a more general proxy for overall actin organization. Indeed, we found that actin fibers in the anterior region of cells displayed significantly higher L_p_ compared to those in the posterior (Fig 3B-D).

**Figure 3:**
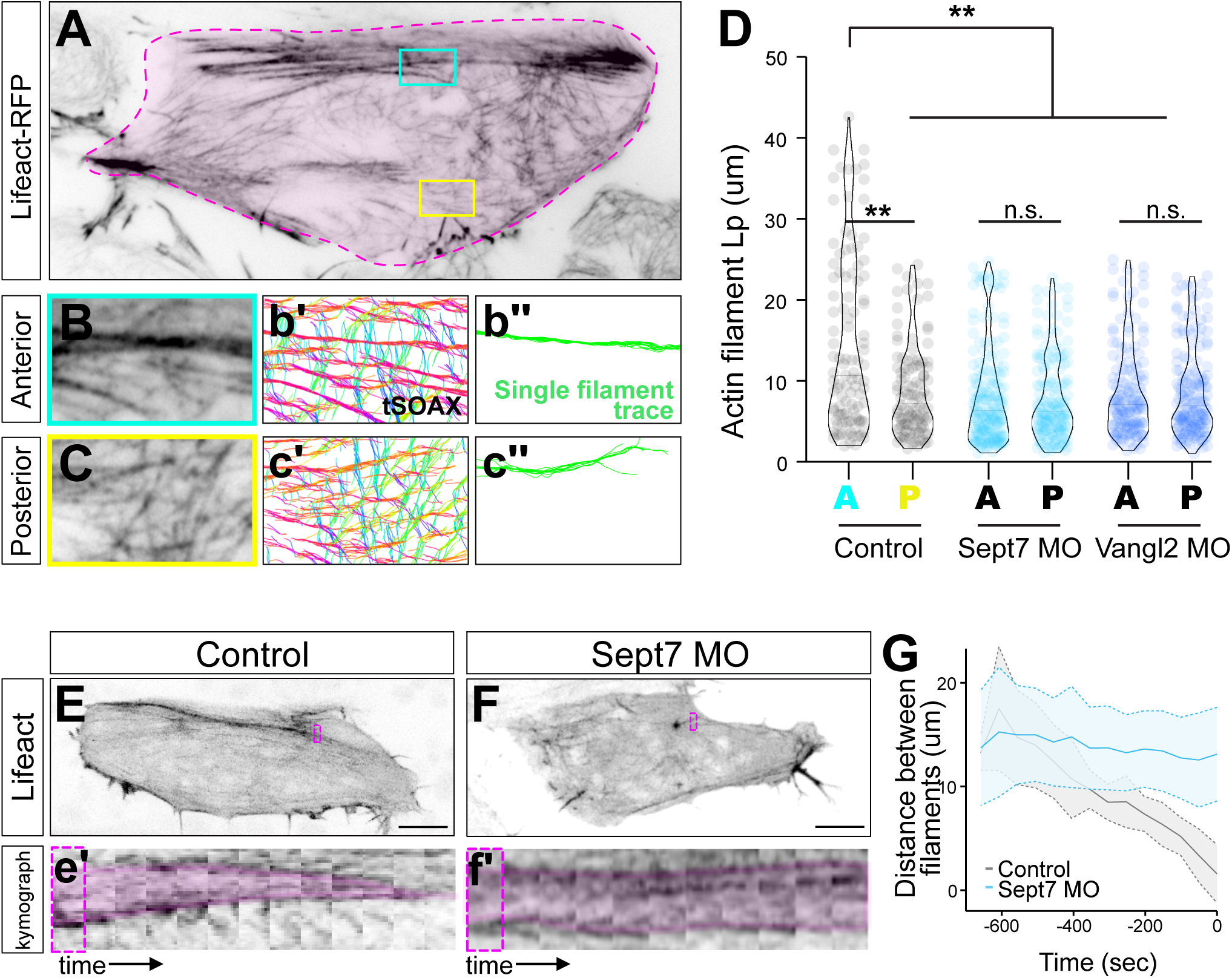
PCP- and Septin-dependent anteroposterior pattern in the node and cable system. **A.** Representative TIRF image of a control cell, outlined in pink. Boxes highlighting anterior (cyan) and posterior (yellow) regions of the cell. **B, C.** Representative anterior and posterior cropped ROIs from A. **b’-c’**. tSOAX segmentation. **b”-c”**. representative single actin filament traces used for quantification. **D**. Quantification of persistence length (Lp) of actin filaments in anterior (A) or posterior (P) cellular regions of control, Sept7MO, and Vangl2MO cells. **E, F.** Lifeact-RFP labeling the actin cortex in control and Sept7 MO cells. Boxes indicate regions used for kymograph, below. **e’, f’.** Kymograph of representative actin fiber coalescence over time. **G.** quantification of actin filament coalescence over time. N values and statistics are provided in Supplemental Information.

Strikingly, the anteroposterior-polarized asymmetry in L_p_ of actin fibers was eliminated by loss of either Vangl2 or Sept7 (Fig. 3D; Supp Fig 5), an effect resulting specifically from reduction in actin persistence lengths in anterior regions (Fig 3D). Together, these findings reveal that the actin node and cable system displays two vectors of polarity: individual actin fibers align mediolaterally in a Vangl2-dependent manner, while higher level actin organization quantified either by cable positioning or by L_p_ of actin fibers is patterned anteroposteriorly in a manner that requires both Vangl2 and Sept7.

### Sept7 is required for progressive coalescence of the anterior actin cables

The data so far indicate that Vangl2 acts via Sept7 to control anteroposterior actin organization in the node and cable system. To avoid the additional pleiotropic effects of core PCP loss, we focused a deeper exploration on Sept7 KD, starting with a time-lapse analysis of the evolution of the Sept7 KD phenotype over time (Fig. 3E, F). In movies, we observed that while individual, smaller actin cables form in the absence of Sept7, they fail to coalesce into a single, anteriorly positioned cable. Using kymographs, we quantified this phenotype, revealing that after Sept7 KD, coalescence was severely disrupted, with actin filaments remaining well-spaced on average, even at later time points (Fig. 3e’, f’, G).

### Sept7 controls local dynamics of polarized actin flow

Finally, we probed the effect of Sept7 loss on actin dynamics using fluorescent speckle microscopy ^29^, in which mosaically labeled actin monomers trace both the bulk flow of actin and the turnover of individual actin filaments ^30,31^. Specifically, we imaged the overall actin network with LifeAct-RFP and identified nodes with Prickle2-GFP, as shown in Fig. 4A, with an enlarged region and split channels shown in Fig. 4B, C. In the same cells, we also imaged single actin speckles using dye-labelled actin monomers (Fig 4A, D).

**Figure 4:**
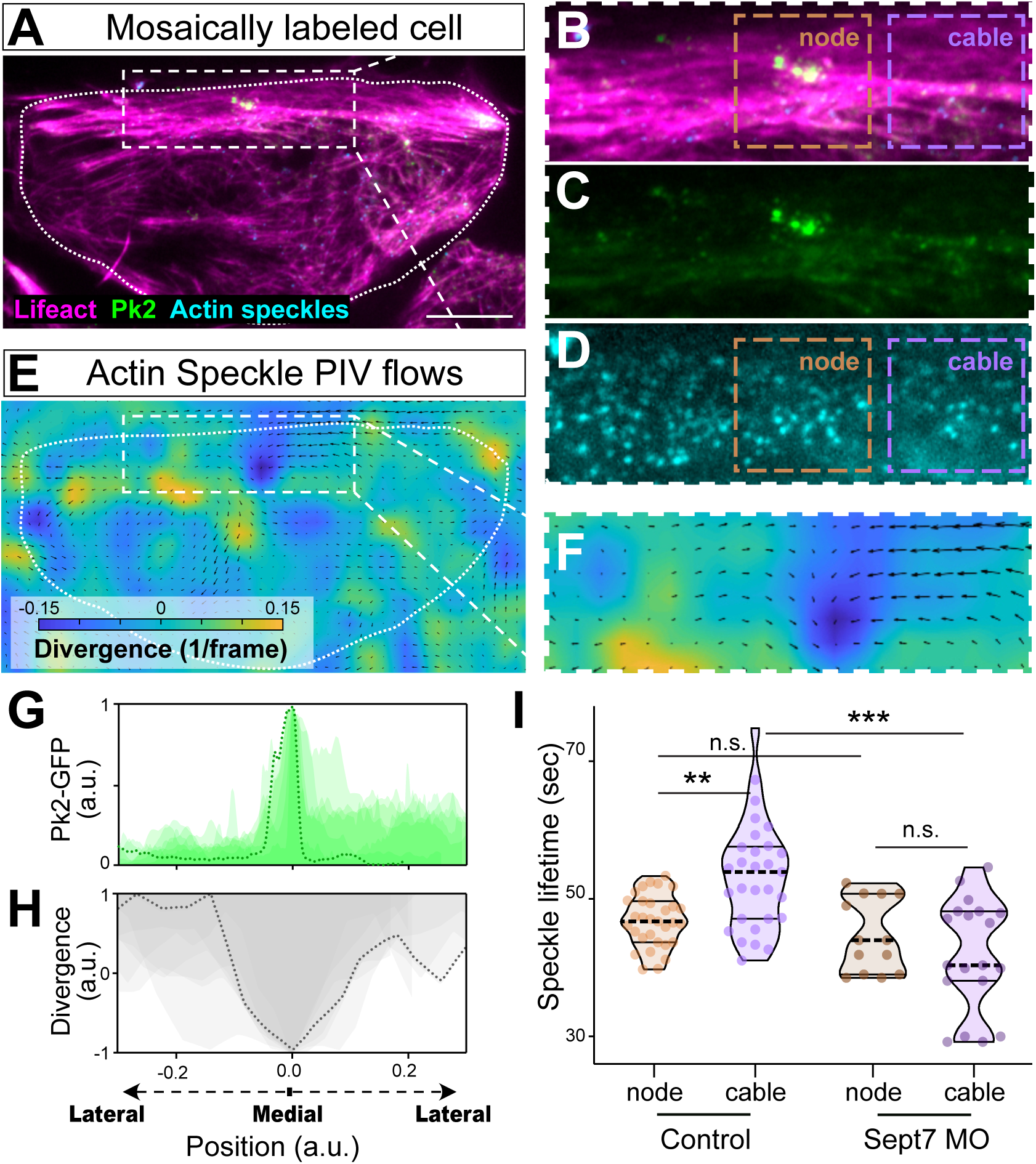
Sept7 controls local dynamics of polarized actin flows. **A.** TIRF image of a single cell labeled with Lifeact-RFP, Pk2-GFP, and actin-647 speckles. **B-D.** Zoomed, split channels of the boxed region in A. Orange and purple boxes indicate regions quantified in Panel I, below). **E.** Heatmap and vector diagram of PIV data for the region shown in A. Arrows indicate the vector and direction of flow; legend shows heatmap indicates positive divergence of flow (blue) or negative divergent (i.e. convergence, yellow). **F.** Zoomed view of the boxed region of the heatmap in E. **G.** Line plot of Pk2-GFP pixel intensities in multiple cells. **H.** Line plot showing vector divergence along the same line from which Pk2 intensity was derived, above. Note maximal vector convergence is coincident with the peak of Pk2 intensity (i.e., in nodes). **I.** Quantification of actin speckle lifetimes in regions indicated by boxes in panels B and D, above. N values and statistics are provided in Supplemental Information.

We then used Particle Imaging Velocimetry (PIV)^32^ analysis to understand the bulk flow of actin. Tracing actin speckles revealed a consistent pattern in which actin flows medially along anterior actin cables into Pk2+ nodes (Fig. 4E, F; arrows indicate the vector and magnitude of particle flow). To quantify flow, we measured the divergence of speckle trajectories over time which we represented with a heat map where yellow represents vector divergence, and blue represents negative divergence (i.e convergence of two vectors). We consistently observed a dark blue region at the anterior of cells (Fig. 4E, F), correlating with the Pk2+ node, indicating that actin speckles converge at the node. To quantify this effect across samples, we overlaid line plots of Pk2-GFP (Fig. 4G) intensity with line plots of vector divergence (Fig. 4H), and these bulk data reveal a peak of Pk2-GFP intensity at nodes coincident with the inverted peak of negative vector divergence (Fig. 4G, H).

Noting that actin frequently flows to regions of enhanced depolymerization and that actin depolymerizing factors are required for normal CE ^4,5,33^, we examined several such factors. We found strong enrichment of Cfl1, Twf1, and Cap in actin nodes (Supp. Fig. 6), suggesting that the node is a region of high actin turnover and that PCP proteins may organize actin flows through the recruitment of such factors.

Finally, we quantified actin filament turnover by tracking *individual* actin speckles over time, independently measuring lifetimes in nodes and in nearby cables (Fig. 4B, D; cyan and purple boxes, respectively). We found that the lifetime of actin speckles in nodes was significantly shorter than the lifetimes in the cable (Fig. 4I, left), suggesting faster turnover at nodes and increased actin filament stability in anterior cables. Because Sept7 is required for cable formation anteriorly (Fig. 3E-G), we asked how loss of Sept7 impacts dynamics of actin speckles in nodes and cables, and we observed a very specific effect: Sept7 KD significantly reduced the longer actin speckle lifetimes observed in cables (Fig. 4I, right), but did not alter the shorter lifetime of actin speckles in nodes (Fig. 4I). These data suggest a role for Sept7 in stabilizing actin in the cable region.

## Conclusions

Here, we exploited the actin node and cable system of *Xenopus* mesoderm cells engaged in CE for an in-depth analysis of the relationship between PCP proteins, Septins, and actin, providing two important insights. First, while the superficial actin node and cable system is essential for CE in *Xenopus* ^16,17^, and its dynamics are controlled by PCP ^18^, how the overt planar polarization of the system relates to PCP proteins has not been obvious. For example, both myosin-driven contraction of the system ^16–18^ and the orientation of actin fibers within it ^5^ are polarized mediolaterally, yet PCP proteins are well-known to mark the anteroposterior axis of cells engaged in CE ^22–24,34,35^. Our findings here are important then for revealing a previously cryptic anteroposterior-directed planar polarity of the actin node and cable system, displayed both by the anterior coalescence of actin cables (Fig. 2) and by the longer persistence length of actin fibers anteriorly (Fig. 3). Crucially, PCP proteins and Sept7 are essential for both.

Second, when considered as a whole, our data suggest a preliminary model for the emergence of both mediolateral and anteroposterior polarity in the node and cable system. PCP proteins (Fig. 2) and Twf1^5^ are each essential for mediolateral alignment of actin fibers, arguing they act in concert and independently of Sept7 to govern this aspect of polarity. At the same time, PCP proteins are anteriorly biased (Fig. 1), where they help to establish the higher-level, anteroposteriorly polarized actin organization described above. Specifically, the order observed anteriorly after loss PCP or Septins is similar to that normally observed posteriorly (Fig. 3). In addition, speckle microscopy revealed that Sept7 is required to stabilize actin filaments in anterior cables. Altogether, these data suggest that PCP may recruit Sept7, where it imparts stability to actin and increases overall order, which in turn facilitates cable formation specifically in anterior regions of cells.

Finally, these data on polarization of an actin network may shed light on our emerging understanding that very local, sub-cellular mechanical heterogeneities play key roles in cell behavior ^36–38^. Both experiments and theory now argue that such heterogeneities are essential for normal CE ^39–41^. It is tempting to speculate that the highly local patterning of the actin cytoskeleton described here may provide a basis for local patterning of cell mechanics.

## Acknowledgements

We would like to acknowledge the Microscopy and Imaging Facility of the Center for Biomedical Research Support at The University of Texas at Austin for use of equipment and specifically thank Anna Webb and Julie Hayes for assistance and technical expertise. And a big thank you to all of the members of the Wallingford lab for their advice and critical feedback!

## Author contributions

C.C.D. and J.B.W. conceived the project. C.C.D., J.B.W., S.W., and J.A. designed the experiments. C.C.D. preformed all imaging and analysis, with assistance in persistence length analysis from S.W. and V.B.P. and J.A. C.C.D and J.B.W. wrote the paper with input and editing from all authors. Funding acquisition from J.B.W.

## Supplemental Methods

### Key resources table

| REAGENT or RESOURCE | SOURCE | IDENTIFIER |
| --- | --- | --- |
| **Antibodies** |  |  |
| Rat anti-Vangl2 antibody (GST-tagged recombinant Vangl2 N-term fragment) | Millipore-Sigma | MABN750 |
| **Chemicals, peptides, and recombinant proteins** |  |  |
| hCG (human chorionic gonadotropin) | Merck | Code# 133754 |
| Alexa 647 conjugated Actin | ThermoFisher | Cat# A12374 |
| **Critical commercial assays** |  |  |
| mMessage mMachine SP6 transcription kit | ThermoFisher | Cat# AM1340 |
| **Oligonucleotides** |  |  |
| Vangl2 morpholino | Park & Moon<sup>55</sup> | Gene-tools.com |
| Pk2 morpholino | Butler & Wallingford<sup>56</sup> | Gene-tools.com |
| Septin7 morpholino | Kim et al<sup>29</sup> | Gene-tools.com |
| **Recombinant DNA** |  |  |
| Lifeact-RFP | Riedl, et al. <sup>57</sup> | N/A |
| Vangl2-GFP | Butler & Wallingford<sup>56</sup> | N/A |
| Pk2-GFP | Butler & Wallingford<sup>56</sup> | N/A |
| Dvl2-GFP | Wallingford, et al.<sup>28</sup> | N/A |
| Septin7-GFP | Becker-Heck, et al.<sup>58</sup>, Shindo & Wallingford<sup>5</sup> | N/A |
| Septin2-GFP | Becker-Heck, et al.<sup>58</sup>, Shindo & Wallingford<sup>5</sup> | N/A |
| Sept8-GFP | Cloned for this paper |  |
| Sept9-GFP | Cloned for this paper |  |
| Twf1-GFP | Devitt, et al.<sup>24</sup> |  |
| Cfl2-GFP | Devitt, et al.<sup>24</sup> |  |
| SF9-mNeon | Vielemeyer, et al.<sup>59</sup> | N/A |
| CAP1-GFP | Devitt, et al.<sup>24</sup> | N/A |
| DN Dishevelled (Xdd1) | Wallingford, et al.<sup>28</sup> | N/A |
| **Software and algorithms** |  |  |
| Prism | GraphPad Software | Graphpad.com |
| Excel | Microsoft | Microsoft.com |
| Matlab | MathWorks | Mathworks.com |
| Python | Python | Python.org |
| Orientation J | Püspöki, et al.<sup>61</sup> | http://bigwww.epfl.ch/demo/orientation/ |
| MosaicSuite | Sbalzarini & Koumoutsakos<sup>62</sup> | http://mosaic.mpi-cbg.de |
| TSOAX v0.1.8 | Xu, et al.<sup>63</sup> | https://www.lehigh.edu/~div206/tsoax/ |

### Lead contact

Further information and reagent requests should be directed to John B. Wallingford.

### Material availability

All reagents available from lead contact with completed MTA upon request.

### Data and code availability

Any additional information required to reanalyze the data reported in this paper is available from the lead contact upon request.

### Data

All data reported in this paper will be shared by the lead contact upon request.

### Code

This paper does not report original code.

### Experimental model and subject details

#### Xenopus embryo manipulations

*Xenopus laevis* females were super-ovulated by injection of hCG (human chorionic gonadotropin). The following day, eggs were squeezed from the females. In vitro fertilization was performed by homogenizing a small part of a harvested testis in 1X Marc’s Modified Ringer’s (MMR) and mixing with collected eggs. Embryos were dejellied in 3% cysteine (pH 7.9) at the two-cell stage. Fertilized embryos were rinsed and reared in 1/3X Marc’s Modified Ringer’s (MMR) solution. For microinjections, embryos were placed in a 2% ficoll in 1/3X MMR solution and injected using forceps and an Oxford universal micromanipulator. Morpholino oligonucleotide or CRISPR/Cas9 was injected into two dorsal blastomeres to target the dorsal marginal zone (DMZ). Embryos were microinjected with mRNA at the 4-cell stage for uniform labeling and at the 64- to 128- cell stage for mosaic labeling. Embryos were staged according to Nieuwkoop and Faber (Nieuwkoop and Faber, 1994). All animal work has been approved by the IACUC of UT, Austin, protocol no. AUP-2021-00167.

### Method details

#### Plasmids, mRNA, protein, and MOs for microinjections

*Xenopus* gene sequences were obtained from Xenbase (www.xenbase.org) and open reading frames (ORF) of genes were amplified from the *Xenopus* cDNA library by polymerase chain reaction (PCR), and then are inserted into a pCS10R vector fused with eGFP. The following constructs were cloned into pCS vector: Sept8-GFP, Sept9-GFP. These constructs were linearized and the capped mRNAs were synthesized using mMESSAGE mMACHINE SP6 transcription kit (ThermoFisher Scientific, Waltham, MA, USA). Concentration for GFP localization was titrated to the lowest concentration where we could still detect GFP signal, but no cellular phenotype was observed. The amount of injected mRNAs per blastomere are as follows: Lifeact-GFP or –RFP [50-100pg], Pk2-GFP [35pg]^6^, Vangl2-GFP [35pg]^6^, Dvl2-GFP [5- 8pg]^28^, SF9-GFP [25pg]^59^, Cap1-GFP[25pg], Septin2- GFP [50pg]^29^, Septin7-GFP [35pg]^5, 58^, Septin8-GFP [40pg], Septin9GFP [40pg] and Xdd1 [500pg]^28^.

#### Morpholinos

Morpholinos used were previously designed to target exon-intron splicing junction (Gene Tools, Philomath, OR, USA). The MO sequences and the working concentrations include: Septin7 morpholino^5, 29, 64^, 5’ TGCTGTAGAGTCAGTGCCTCGCCTT 3’, 40 ng per injection at 4-cell stage; Vangl2 morpholino^55^, 5’ CGTTGGCGGATTTGGGTCCCCCCGA 3’, 20ng per injection at 4-cell stage; Pk2 morpholino^56^, 5’ GAACCCAAACAAACACTTACCTGTT 3’, 20ng per injection at 4-cell stage.

#### Actin Speckle Microscopy

For fluorescent actin speckle microscopy, labeled actin monomers (Alexa 647 conjugated Actin, Catalog number A12374, ThermoFisher, Waltham, MA, USA; initial experiments were performed with Alexa 568, yielding the same results) were reconstituted in a 2mg/ml stock solution. A 0.01ug/ul working solution was prepared for each 10nl injection, and 0.1ng was injected into dorsal blastomeres of 4-cell stage embryos. This concentration was determined empirically by titrating the amount injected until no cellular phenotype was observed and labeled actin monomers were sparse enough to detect individual “speckles” ^40^.

#### Live imaging of Keller explants

mRNAs were injected at the dorsal side of 4 cell stage embryos, and the dorsal marginal zone (DMZ) tissues were dissected out at stage 10.5 using forceps, hairloops, and eyebrow hair knives. Each explant was mounted to a fibronectin-coated dish in Steinberg’s solution^5^, and cultured at 16℃ for half a day before imaging with a Zeiss LSM700 confocal microscope (Jena, Germany), or Nikon N-STORM combined TIRF/STORM microscope (Minato City, Tokyo, Japan).

#### Inhibitor treatments

Inhibitors were solubilized in DMSO to a stock concentration of 10mM for Latrunculin B and 100mM for (+) Blebbistatin. Explants with labeled actin and Vangl2 were treated with 1uM LatrunculinB (diluted 1:10,000 in Steinberg’s solution)(CAS Number: 76343-94-7), a cell permeable actin polymerization inhibitor, or 100uM (+) Blebbistatin (diluted 1:1,000 in Steinberg’s solution) (CAS Number 856925-75-2), a non-muscle myosin inhibitor. Samples were treated with DMSO (vehicle control) or specific inhibitors and imaged immediately after application.

#### Image analysis

Images were processed with the Fiji distribution of ImageJ and Photoshop (Adobe) software suites, and figures were assembled in Illustrator (Adobe) and Affinity Designer (***).

*Cell length and width measurements* were manually taken in Fiji along the long axis and through the centroid of the cell.

*AP positioning of node and PCP proteins* measurements were taken by manually drawing a vertical line though the AP axis of a cell through the node and cable region. Intensity plots were measured using the multiplot feature in Fiji. The pixels of highest fluorescent intensity were identified for proteins of interest and the corresponding position within the cell was noted. To compare across many samples, this position was scaled between 0 and 1 with 0 representing the most anterior position within a cell and 1 representing the most posterior position.

Similarly, *Anterior cable intensity measurements* were taken by drawing a vertical line through the midline of the cell with a line width matching the average width of a single cell (150 pixels). Using the multiplot feature in Fiji, the average actin intensity averaged across the anterior posterior axis of a cell was quantified and averaged over several samples.

*Actin cable orientations* were taken from TIRF images of single cell cortical actin network using the OrientationJ plugin in Fiji ^61^ (http://bigwww.epfl.ch/demo/orientation/).

*Actin cable bundling measurements* were taken using kymographs and manual measurements between two points in Fiji. Kymographs were taken across the anteroposterior axis of individual cells near the actin node. We then measured the distance between adjacent actin filaments over time over specified time intervals.

*Divergence measurements of actin speckles* were taken using the PIVlab^42^ software in Matlab. Specific xy coordinates within the cell were identified to link divergence measurements obtained in PIVlab/Matlab with intensity values demarcating the node taken with Fiji.

*Fluorescent actin speckle lifetime* measurements were taken by tracking individual speckles using MosaicSuite^62^ (http://mosaic.mpi-cbg.de) object tracking plugin in Fiji. Custom scripts were written in Matlab to count the lifetime of each speckle. We defined the node by localization of anterior PCP proteins (Vangl2-GFP or PK2 GPF) or myosin (SF9-GFP) along with focal enrichment of actin. We defined the cable region by actin fiber morphology and localization directly adjacent to the node. Each anterior and posterior actin speckle ROI is 100px x 100px.

#### Persistence length measurements of actin filaments

The open-source biopolymer tracking software TSOAX v0.1.8^63^ (https://www.lehigh.edu/~div206/tsoax/) was used to track actin fibers from TIRF movies automatically. The ridge threshold, stretch-factor, and gaussian std (pixels) parameters were set to 0.01, 1.5, and 1 respectively. These values were chosen as they resulted in accurate filament labeling. The remaining parameters were unchanged from default values. TIRF movies were processed prior to TSOAX analysis with the application of a gaussian blur of sigma radius = 1 to enhance the accuracy of fiber tracking. TSOAX output containing the spatial and temporal coordinates that define labeled fibers and track groupings was further analyzed to quantify the persistent length of each fiber using customized Python and MATLAB scripts. Briefly, we treated each actin filament as a non-equilibrated semi-flexible polymer. Its persistent length *P* can be obtained via an equation derived from the worm-like chain model (WLC) (cite [49]):

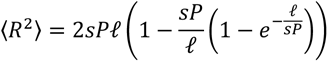

where ℓ is the length of an arbitrary segment of the filament, *R* is the end-to-end distance of the segment, and *s* is set to 1.5 for non-equilibrated fibers. The measure of the mean square of *R* is a function of ℓ for each fiber, and the weighted least squares is used to measure.

#### Quantification and statistical analysis

Statistical analyses were carried out using Microsoft Excel, Matlab (Natick, MA, USA), and GraphPad Prism (Graphpad, San Diego, CA, USA) software. Each experiment was conducted on multiple days and included biological replicates. Due to non-normality observed and low n-number in some observations, we used non-parametric tests for statistical comparisons throughout the manuscript. We used Mann Whitley U tests, Kruskal-Wallis tests, or Kolmogorov–Smirnov tests for statistical comparisons of two groups, multiple groups, or distributions, respectively. MEAN and SD are plotted along with individual data points throughout the manuscript.

## N values and Statistics

**Fig 1:**

*Panel G:* Polarized: Kruskal-Wallis test; Vangl2 vs Pk2 p>0.999, Vangl2 vs Dvl2 p<0.0001, Pk2 vs Dvl2 p<0.0001; Vangl2 GFP n= 74 cells from 10 embryos; Pk2GFP n= 45 cells from 3 embryos; Dvl2-GFP n= 75 cells from 9 embryos)

*Panel H:* Pre-polarized: Kruskal-Wallis, p>0.999 n.s., Vangl2 GFP n= 31 cells from 7 embryos; Pk2GFP n= 42 cells from 9 embryos; Dvl2-GFP n= 84 cells from 3 embryos).

*Panel P:* Mann-Whitney test, control vs Sept7 MO, p<0.0001****, n=34 control cells from 3 embryos; n= 60 Sept7MO cells from 6 embryos).

**Fig. 2:**

*Panels A-F:* Hr1 n= 103 cells, Hr2 n=117 cells, Hr3 n=98 cells, Hr4 n=73 cells).

*Panels G-K:* Vangl 2 MO: Kolmogorov-Smirnov test, actin filament orientation p<0.0001; n= 38 control cells from 12 embryos; n=46 morphant cells from 13 embryos; Pk2MO, KS test, actin filament orientation p<0.0001; n= 23 control cells from 4 embryos; n= 23 morphant cells from 4 embryos). D) dominant negative (DN) disheveled (Xdd1) (control vs Xdd1, Kolmogorov-Smirnov test, actin filament orientation p<0.0001; n=23 control cells from 13 embryos; n= 38 mutant cells from 8 embryos) and E) Sept7 morphant cell, inlay of actin node and cable, magenta box highlighting region of inlay (control vs Sept7MO, Kolmogorov-Smirnov test, actin filament orientation p=0.5596; n= 26 control cells from 10 embryos; n= 31 morphant cells from 10 embryos).

**Fig. 3.**

*Panel D.* Kruskal-Wallis test, (Control A vs P p=0.0046**, Sept7MO A vs P p>0.999 n.s., Vangl2 MO A vs P p>0.999 n.s., Control A vs Sept7MO A p=0.0481*, Control A vs Sept7MO P p= 0.0037**, all other pairwise comparisons p>0.999 n.s.)

*Panel G.* (control vs vangl2 MO, n=16 control cells from 6 embryos; n= 19 Vangl2MO cells from 6 embryos). Scale= 10um.

**Fig. 4.**

*Panel G, H:* n=7 control cells from 6 embryos.

*Panel I:* (Kruskal-Wallis test, control node vs control cable p=0.0093, control node vs Sept7MO node ns p>0.9999, control cable vs Sept7MO cable p<0.0001, Sept7MO node vs Sept7MO cable p>0.999, control cable vs Sept7MO nodes p=0.0033, nodes vs Sept7MO cable n.s. p=0.5498; n= 31 control node, n=29 control cable from 12 embryos; n= 13 Sept7MO node, n= 19 Sept7MO cable from 7 embryos). E) actin cable bundling in control cells.

---

**Figure S1:**
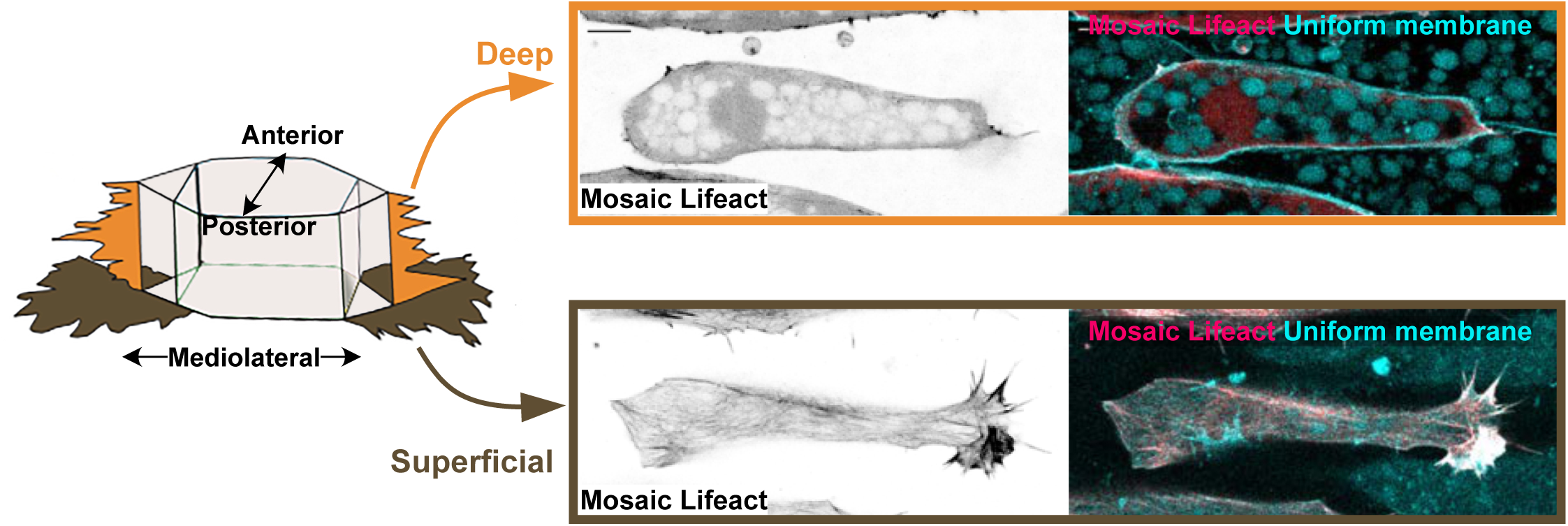
Three distinct actin populations contribute to CE in *Xenopus* gastrula dorsal mesoderm cells. Schematic at left shows a cell in the DMZ with axes labelled, orange and brown structures represent mediloaterally oriented lamellipodia at the superficial and deep level, respectively. Images at right show optical sections through different z-levels of a representative DMZ cell. Actin is labeled by mosaic expression of LifeAct, note anteriorly biased superficial actin cable.

**Figure S2:**
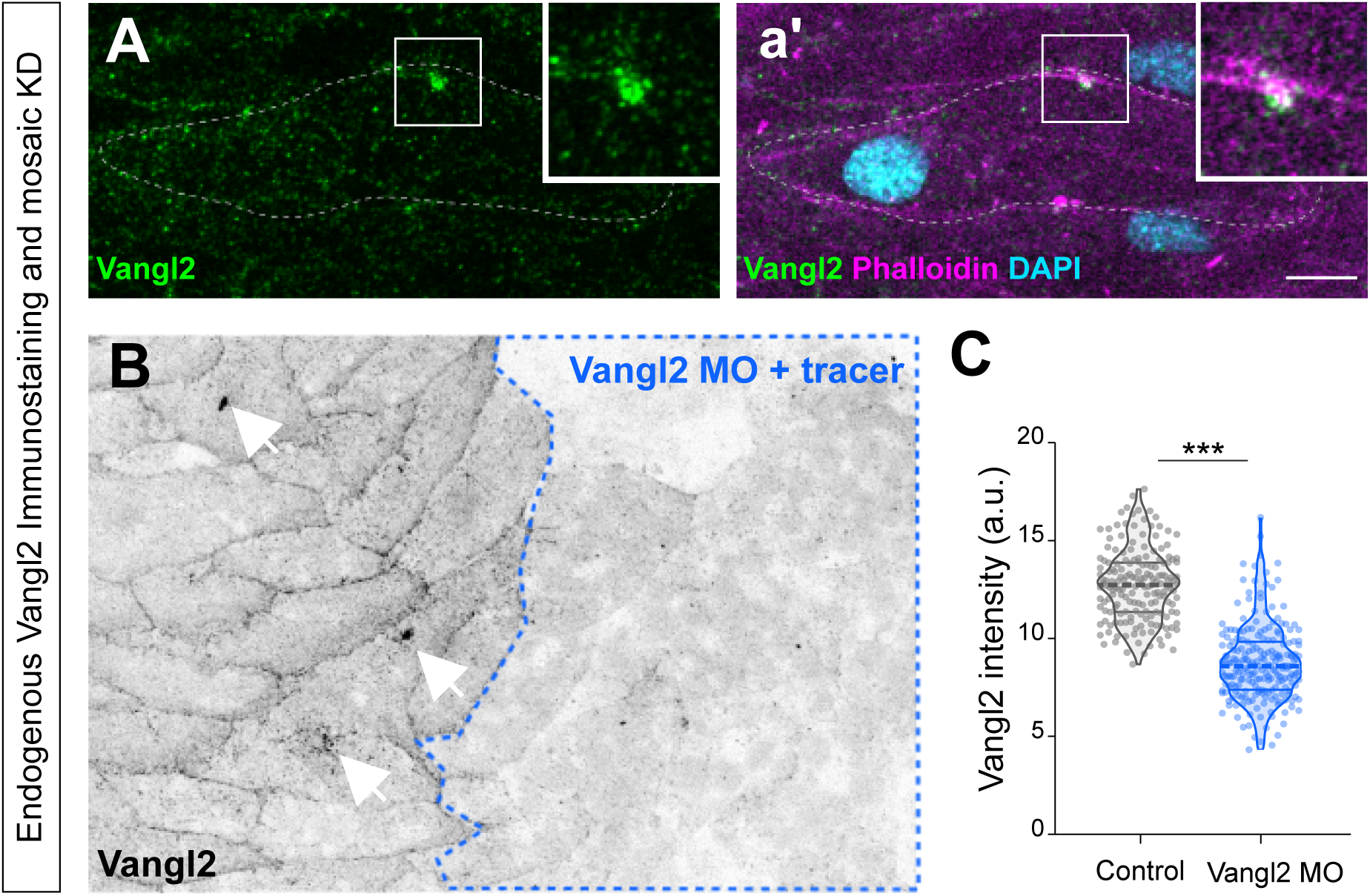
Endogenous Vangl2 localizes to actin-rich nodes. **A.** Immunostaining of Vangl2 (green) along with a’) phalloidin (magenta), and DAPI (cyan) showing co-localization of endogenous Vangl2 with actin-rich nodes. **B.** Endogenous Vangl2 in tissue with mosaic morpholino-mediated KD of Vangl2 (outlined in blue), arrowhead pointing to Vangl2 puncta on control side which are absent in the KD condition. **C.** Quantification confirming mosaic knockdown of Vangl2. N=177 control cells from 3 embryos; n= 219 Vangl2 MO cells from 3 embryos, compared by t test.

**Figure S3:**
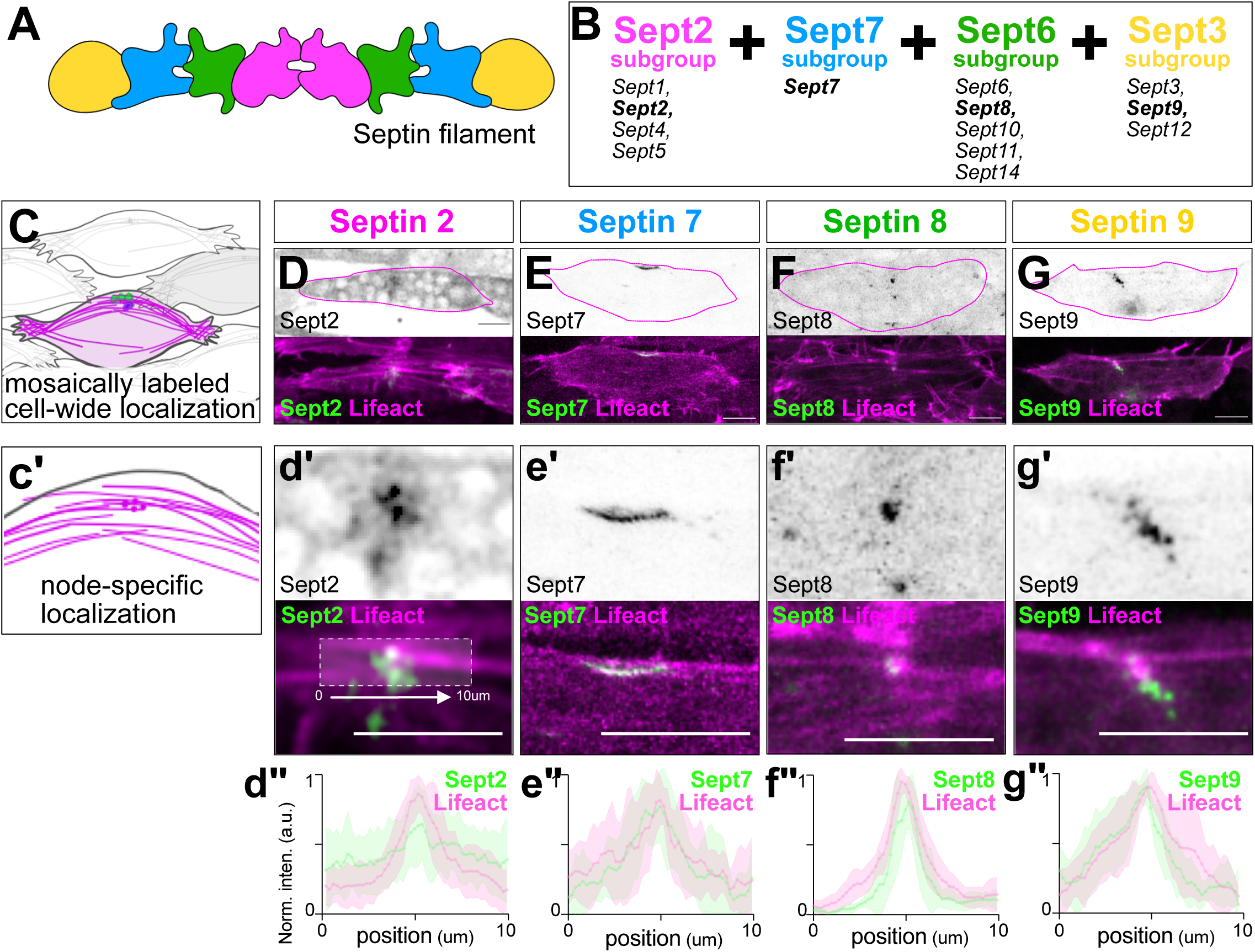
Septins localize to actin-rich nodes. **A.** Schematic of a septin heteroctomer. **B.** Table of combinatorial septin subtypes. **C.** Schematic of representative cell reflecting images in panels D-G. **D-G.** Representative images of localization of indicated GFP-fused septins; merged view of septins (green) and LifeAct are shown below. **c’.** Schematic of zoomed view reflecting images in I-L**. d’-g’.** Zoomed view of indicated septins at actin-rich nodes; merged view of septins (green) and Lifeact are shown below. **d”-g”.** Line intensity plots of Septin GFP fusions and Lifeact intensity across multiple samples. Scale bars=10um.

**Figure S4:**
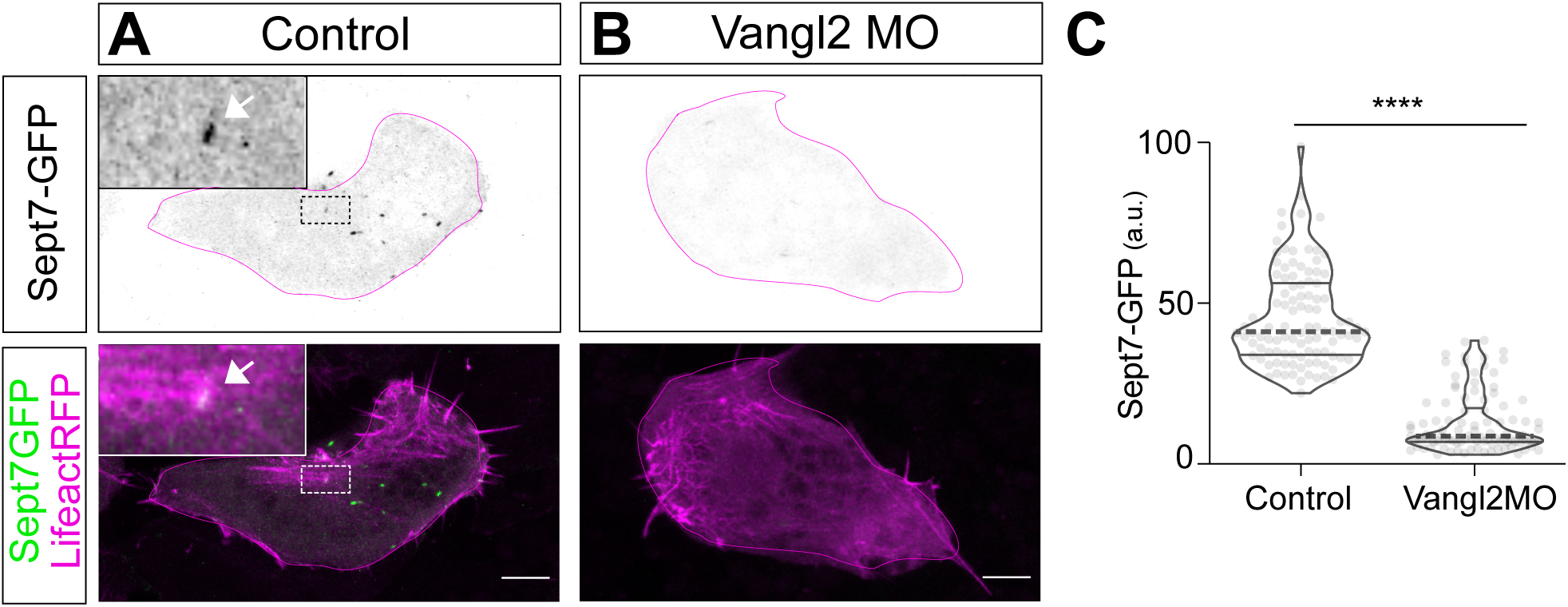
Effect of Vangl2 loss on Sept7. **A.** Representative image of Sept7-GFP in a control cell; merged view of sept7 (green) and Lifeact (magenta) is shown below. **B.** Representative image of Sept7-GFP in a Vangl2 KD cell; merged view of sept7 (green) and Lifeact (magenta) is shown below. **C.** Quantification of Sept7-GFP pixel intensity in control and Vangl2 MO cells. N=102 control cells from 7 embryos; n= 101 Vangl2 MO cells from 6 embryos, compared by t test.

**Figure S5:**
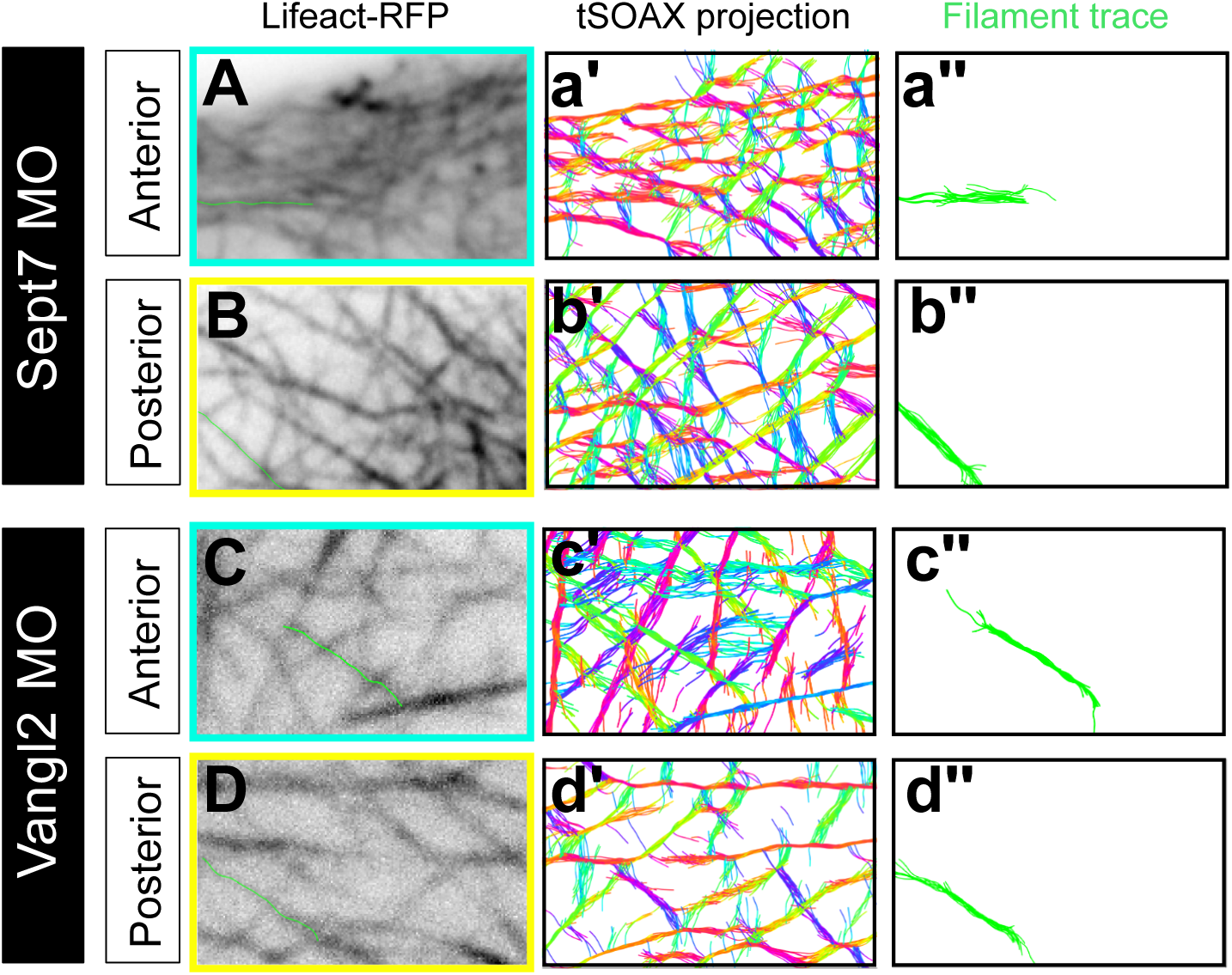
TIRF and SOAX images for Sept7 and Vangl2 MO cells. **A-D.** Representative anterior and posterior cropped regions of interest from TIF movies of Septin7MO or Vangl2MO cells as indicated. **a’-d’**. tSOAX segmentation. **a”-d”**. Representative single actin filament traces used for quantification.

**Figure S6:**
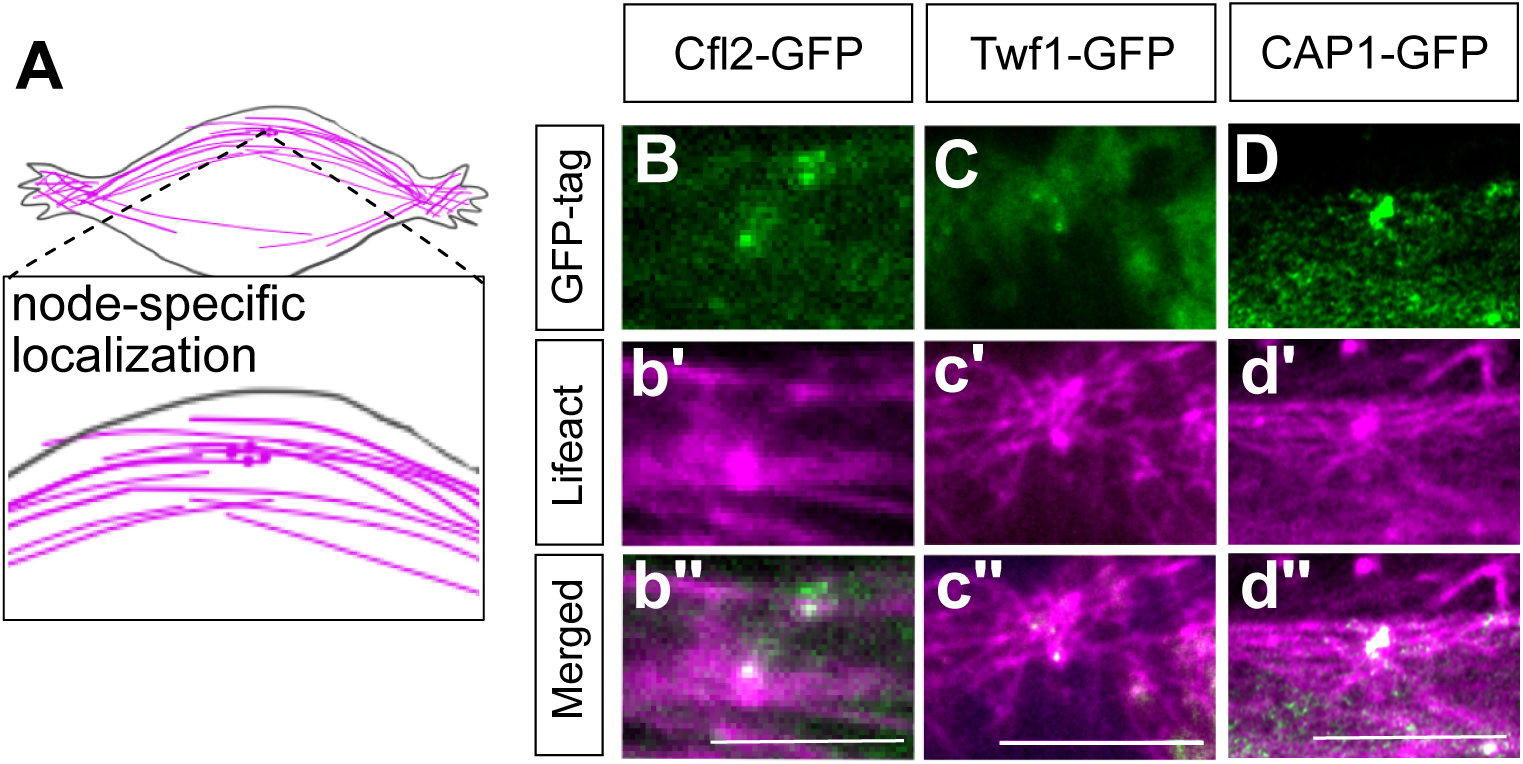
Localization of proteins associated with actin depolymerization. **A.** Schematic of representative cell indicating regions shown images in panels B-D. **B-D.** Representative images of localization of indicated GFP-fused proteins; merged view of the proteins (green) and Lifeact (magenta) are shown below.

